# Mechanistic basis for the evolution of chalcone synthase catalytic cysteine reactivity in land plants

**DOI:** 10.1101/429795

**Authors:** Geoffrey Liou, Ying-Chih Chiang, Yi Wang, Jing-Ke Weng

## Abstract

Flavonoids are important polyphenolic natural products, ubiquitous in land plants, that play diverse functions in plants’ survival in their ecological niches, including UV protection, pigmentation for attracting pollinators, symbiotic nitrogen fixation, and defense against herbivores. Chalcone synthase (CHS) catalyzes the first committed step in plant flavonoid biosynthesis and is highly conserved in all land plants. In several previously reported crystal structures of flowering plant CHSs, the catalytic cysteine is oxidized to sulfinic acid, indicating enhanced nucleophilicity in this residue associated with its increased susceptibility to oxidation. In this study, we report a set of new crystal structures of CHSs representing all five major lineages of land plants. We reveal that the structures of CHS from a lycophyte and a moss species preserve the catalytic cysteine in a reduced state, in contrast to the cysteine sulfinic acid seen in all euphyllophyte CHS structures. *In vivo* complementation, *in vitro* biochemical and mutagenesis analyses, as well as molecular dynamics simulations identify a set of residues that differ between basal-plant and euphyllophyte CHSs and modulate catalytic cysteine reactivity. We propose that the CHS active-site environment has evolved in euphyllophytes to further enhance the nucleophilicity of the catalytic cysteine since the divergence of euphyllophytes from other vascular plant lineages 400 million years ago. These changes in CHS could have contributed to the diversification of flavonoid biosynthesis in euphyllophytes, which in turn contributed to their dominance in terrestrial ecosystems.

## Introduction

In the transition from aquatic domains to terrestrial environments, early land plants faced several major challenges, including exposure to damaging UV-B radiation once screened by aquatic environments, lack of structural support once provided by buoyancy in water, drought, and co-evolving herbivores and pathogens. To cope with many of these stresses, land plants have evolved a series of specialized metabolic pathways, among which phenylpropanoid metabolism was probably one of the most critical soon after the transition from water to land (Weng and Chapple, 2010).

Flavonoids are a diverse class of plant phenolic compounds found in all extant land plants, with important roles in many aspects of plant life, including UV protection, pigmentation for attracting pollinators and seed dispersers, defense, and signaling between plants and microbes (Winkel-Shirley, 2001). Some flavonoids are also of great interest for their anti-cancer and antioxidant activities as well as other potential health benefits to humans (Yao et al., 2004). After the core flavonoid biosynthetic pathway was established in early land plants, new branches of the pathway continued to evolve over the history of plant evolution, producing structurally and functionally diverse flavonoids to cope with changing habitats, co-evolving pathogens and herbivores, and other aspects of plants’ ecological niches. Basal bryophytes biosynthesize the three main classes of flavonoids, namely flavanones, flavones, and flavonols, which likely emerged as UV sunscreens (Rausher, 2006). The lycophyte *Selaginella* biosynthesizes a rich diversity of biflavonoids, many of which were shown to be cytotoxic and may function as phytoalexins (Weng and Noel, 2013). The ability to synthesize the astringent, polyphenolic tannins, which defend against bacterial and fungal pathogens, seems to have evolved in euphyllophytes (Rausher, 2006). Finally, seed plants, including gymnosperms and angiosperms, developed elaborate anthocyanin biosynthetic pathways to produce the vivid colors used to attract pollinators or ward off herbivores.

Chalcone synthase (CHS), a highly conserved plant type III polyketide synthase (PKS), is the first committed enzyme in the plant flavonoid biosynthetic pathway. CHS synthesizes naringenin chalcone from a molecule of *p*-coumaroyl-CoA and three molecules of malonyl-CoA (Weng and Noel, 2012) (Figure 1A). The proposed catalytic mechanism of CHS involves loading of the starter molecule *p-* coumaroyl CoA onto the catalytic cysteine, which also serves as the attachment site of the growing polyketide chain during the iterative elongation steps (Austin and Noel, 2003). This initial reaction step requires the cysteine to be present as a thiolate anion before loading of the starter molecule (Figure 1B). Using thiol-specific inactivation and the pH dependence of the malonyl-CoA decarboxylation reaction, the p*K*_a_ of the catalytic cysteine (Cys 164) of *Medicago sativa* CHS (MsCHS) was measured to be 5.5, a value significantly lower than 8.7 for free cysteine (Jez and Noel, 2000).

**Figure 1.**
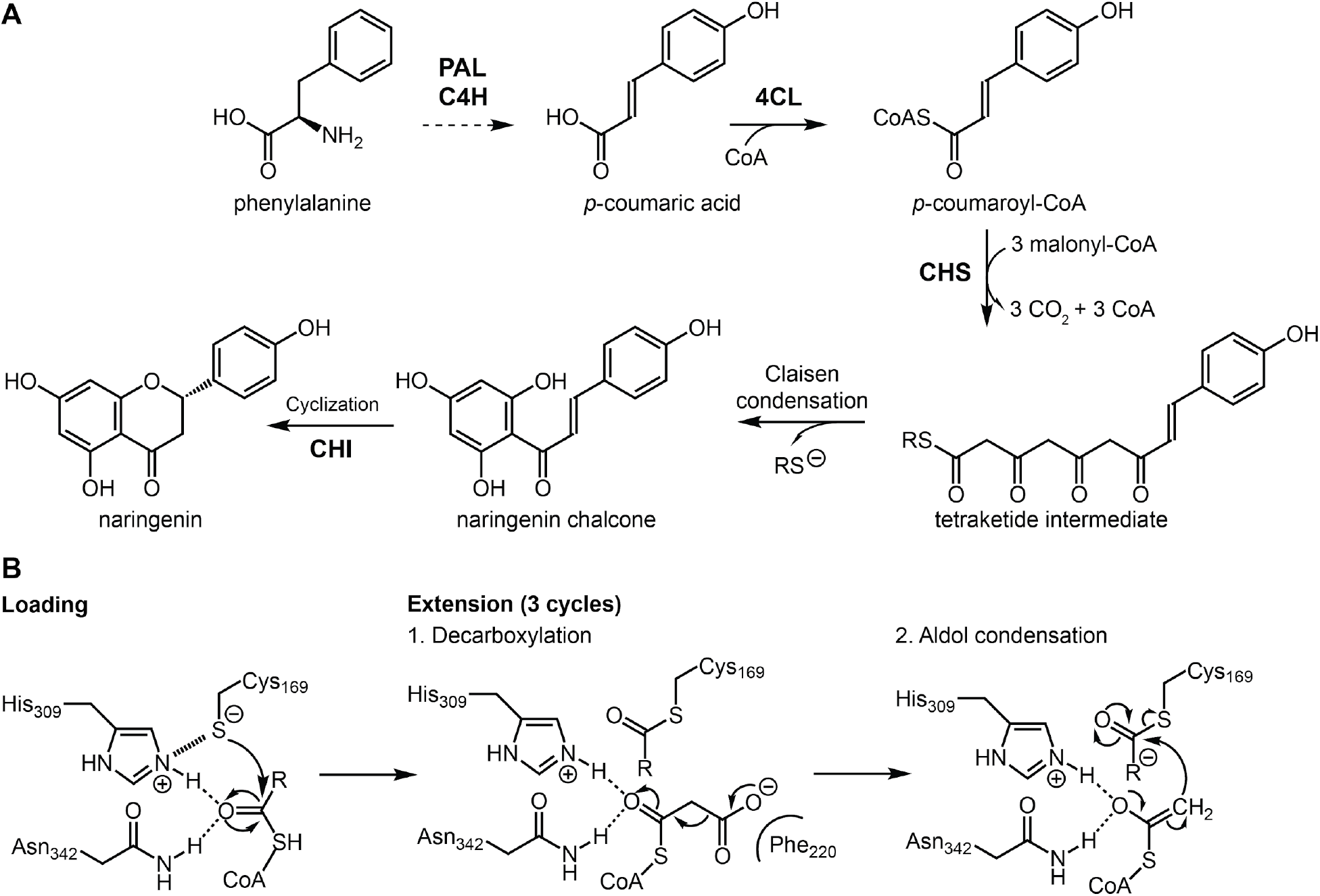
**A**, Phenylpropanoid and flavonoid metabolism. PAL, phenylalanine ammonia-lyase; C4H, *trans*-cinnamate 4-monooxygenase; 4CL, 4-coumarate-CoA ligase; CHS, chalcone synthase; CHI, chalcone isomerase; CoA, coenzyme A. Cyclization of naringenin chalcone to naringenin also proceeds spontaneous in aqueous solution. **B**, Reaction mechanism of CHS. The extension step is performed three times to repeatedly extend the starter molecule malonyl-CoA to form a linear tetraketide intermediate, which then cyclizes to form naringenin chalcone.

Interestingly, we observed that the catalytic cysteine residues in the previously reported MsCHS crystal structures appear to be oxidized to sulfinic acid (Ferrer et al. 1999). Furthermore, the same phenomenon was observed in the crystal structures for several other plant type III PKSs evolutionarily derived from CHS, including *Gerbera hybrida* 2-pyrone synthase (Jez et al., 2000) (Supplementary Figure 1). The other non-catalytic cysteines in these proteins do not appear to be oxidized. These findings suggest that the oxidation of the catalytic cysteine observed in several type III PKS crystal structures may not simply be an artifact of X-ray crystallography, but rather reflects the intrinsic redox potential and the reactivity of the catalytic cysteine evolved in this family of enzymes. Indeed, the propensity for a particular cysteine residue to undergo oxidation has been previously indicated to correlate with low p*K*_a_ (Reddie and Carroll, 2008).

Here, we present a set of new crystal structures of orthologous CHSs representing five major lineages of land plants, namely bryophytes, lycophytes, monilophytes, gymnosperms, and angiosperms, spanning 500 million years of land plant evolution. Through comparative structural analysis, *in vivo* complementation, *in vitro* biochemistry, mutagenesis studies, and molecular dynamics simulations, we reveal that CHSs of basal land plants, i.e. bryophytes and lycophytes, contain a catalytic cysteine less reactive than that of the CHSs from higher plants, i.e. euphyllophytes. We probe into the structure-function relationship of a set of residues that modulate the nucleophilicity of the catalytic cysteine, which leads us to propose that euphyllophytes may have evolved a more catalytically efficient CHS to enhance flavonoid biosynthesis relative to their basal plant relatives.

## Results

### Basal-plant CHSs contain reduced catalytic cysteine in their crystal structures

To examine the structural basis for the evolution of CHS across major land plant lineages, we cloned, expressed, and solved the crystal structures of the five CHS orthologs from the moss *Physcomitrella patens* (PpCHS), the lycophyte *Selaginella moellendorffii* (SmCHS), the monilophyte *Equisetum arvense* (EaCHS), the gymnosperm *Pinus sylvestris* (PsCHS), and the angiosperm *Arabidopsis thaliana* (AtCHS) (Figure 2A and B, Supplementary Table 1). Like previously reported crystal structures of type III polyketide synthases, all five CHS orthologs form symmetric homodimers and share the same αβαβα thiolase fold, suggesting a common evolutionary origin (Ferrer et al., 1999). The catalytic triad of cysteine, histidine, and asparagine is found in a highly similar conformation to other PKS and related fatty acid biosynthetic β-ketoacyl-(acyl-carrier-protein) synthase III (KAS III) enzymes, suggesting that they share a similar general catalytic mechanism (Figure 2B).

**Figure 2.**
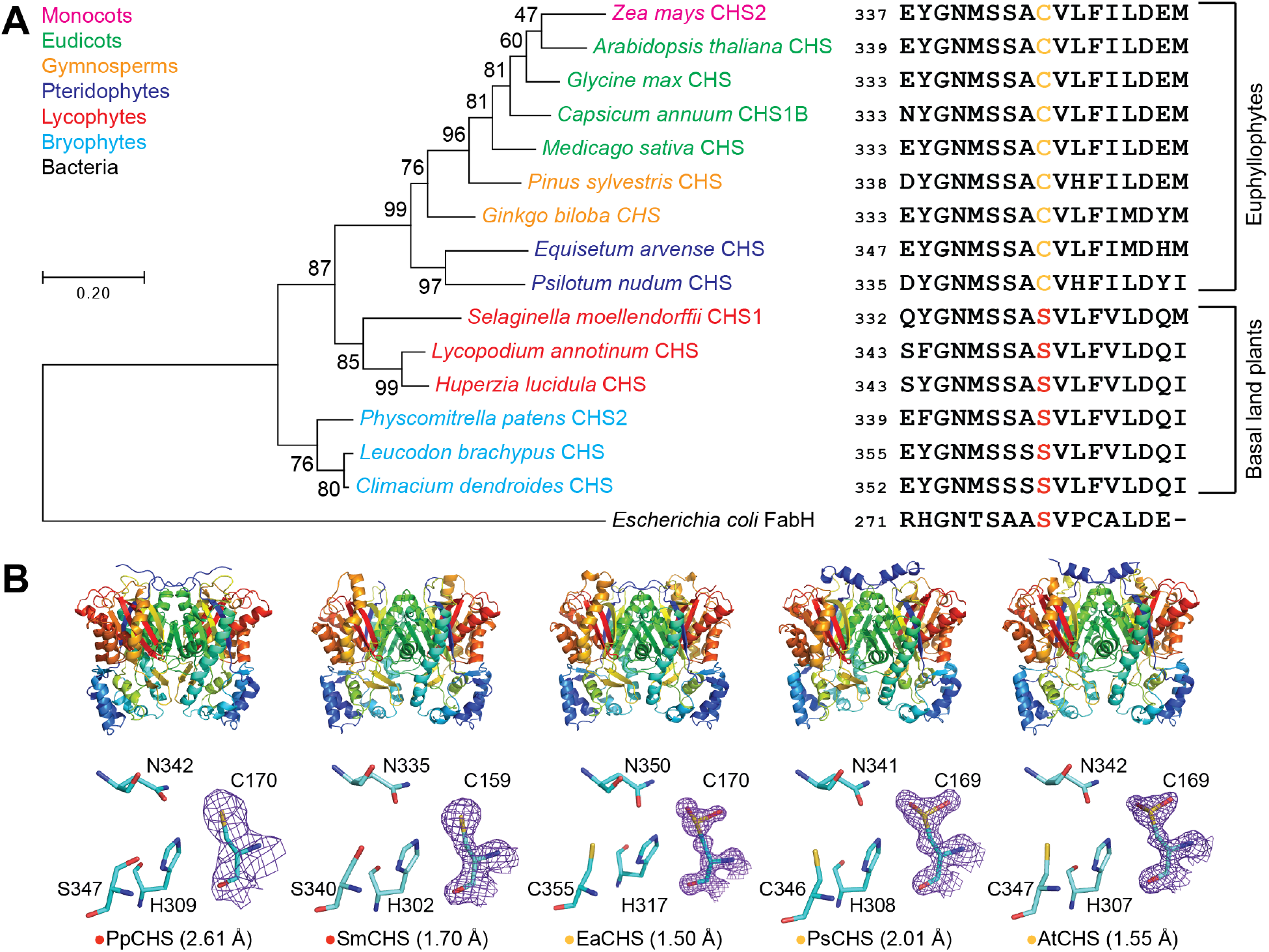
Structural and *in vivo* functional characterization of diverse CHS orthologs. **A**, A maximum-likelihood phylogenetic tree of CHSs from diverse land plant species, with clades indicated by color. The tree is rooted on a bacterial KAS III enzyme (EcFabH). The scale bar indicates evolutionary distance in substitutions per amino acid. **B**, Overall apo crystal structures and active site structures of CHSs from diverse plant lineages. The homodimeric form of CHS is shown with a color gradient from blue at the N terminus to red at the C terminus of each monomer. The 2F_o_-F_c_ electron density map contoured at 1.5σ is shown around the catalytic cysteine. CHSs from euphyllophytes show the catalytic cysteine oxidized to sulfinic acid, whereas CHSs from basal land plants have a reduced catalytic cysteine. The red or yellow dot next to the enzyme name indicates the presence of serine or cysteine, respectively, in position 347 (AtCHS numbering).

Based on the previously proposed reaction mechanism for MsCHS, the catalytic cysteine is C169 in AtCHS and C159 in SmCHS. This residue initiates the reaction mechanism by performing nucleophilic attack on *p*-coumaroyl-CoA (Figure 1B). The other two members of the catalytic triad consist of H309 and N342 in AtCHS, and H302 and N335 in SmCHS. The catalytic histidine contributes to the lowered p*K*_a_ of the catalytic cysteine by forming a stable imidazolium-thiolate ion pair (Jez and Noel, 2000). The histidine and asparagine also form the oxyanion hole that stabilizes the tetrahedral transition states formed during the initial nucleophilic attack by cysteine on *p*-coumaroyl-CoA and after malonyl-CoA decarboxylation (Figure 1B).

Notably, SmCHS and PpCHS are the first CHSs for which a reduced catalytic cysteine has been observed in the crystal structure (Figure 2B). The catalytic cysteine in SmCHS can still become oxidized to sulfenic acid when the crystal is soaked in hydrogen peroxide, indicating that it is still susceptible to oxidation at a lower rate (Supplementary Figure 2). Like most other euphyllophyte type III PKS crystal structures solved to date, AtCHS, PsCHS, and EaCHS contain doubly oxidized catalytic cysteine sulfinic acid (Figure 2B). This interesting observation suggests a functional divide between basal-plant and euphyllophyte CHSs. Despite shared orthology, the redox potential of the catalytic cysteine in PpCHS and SmCHS may differ from that of the euphyllophyte CHSs, resulting in different levels of sensitivity to oxidation under similar crystallization conditions. This could be due to the evolution of some novel molecular features in euphyllophyte CHSs not present in the lower-plant CHSs.

### Basal-plant CHSs only partially complement the Arabidopsis *CHS*-null mutant

CHS orthologs have been identified in all land plant species sequenced to date, suggesting a highly conserved biochemical function. To test whether the five CHSs from the five major plant lineages are functionally equivalent, we generated transgenic *Arabidopsis thaliana* lines expressing each of the five different *CHS*s driven by the Arabidopsis *CHS* promoter in the *CHS*-null mutant *transparent testa 4–2* (*tt4–2*) background (Shirley et al., 1995) (Supplementary Figure 3).

Twenty independent T1 plants were selected for each construct. The phenotypes of the transgenic plants described below were represented by the majority of independent transgenic events for each unique construct. As the name indicates, the *tt4–2* mutant is devoid of flavonoid biosynthesis and therefore lacks the accumulation of the brown condensed tannin pigments in seat coats, revealing the pale yellow color of the underlying cotyledons (Shirley et al., 1995). Whereas At*CHS*, Ps*CHS* and Ea*CHS* fully complement the *tt* phenotype of *tt4–2*, Pp*CHS* and Sm*CHS* only partially rescue the seed *tt* phenotype of *tt4–2* (Supplementary Figure 3), suggesting that Pp*CHS* and Sm*CHS* are likely less active than their higher-plant counterparts *in vivo*. This result also correlates with the crystallographic observation where the catalytic cysteine of basal plant and euphyllophyte CHSs exhibit differential susceptibility to oxidation.

### The p*K*_a_ of the catalytic cysteine is higher in basal-plant CHSs than in euphyllophyte CHSs

To perform nucleophilic attack on the *p*-coumaroyl-CoA substrate, the catalytic cysteine must be present in the thiolate anion form. As shown previously in MsCHS, the p*K*_a_ of the catalytic cysteine is lowered to 5.5, well below physiological pH, in order to stabilize this deprotonated state (Jez and Noel, 2000). Two factors could contribute to the depressed p*K*_a_ of C164. First, H303, one of the catalytic triad of CHS in vicinity of C164, provides an ionic interaction with C164 that can further stabilize the cysteine thiolate anion. Second, C164 is positioned at the N-terminus of the MsCHS α-9 helix (Ferrer et al., 1999), which provides a stabilizing effect on the cysteine thiolate anion through the partial positive charge of the helix dipole (Kortemme and Creighton, 1995). The acidic p*K*_a_ of the catalytic cysteine in CHS ensures the presence of a cysteine thiolate anion in the enzyme active site at physiological pH to serve as the nucleophile for starter molecule loading.

To measure the p*K*_a_ of the catalytic cysteine in the five land plant CHS orthologs, we performed pH-dependent inactivation of CHS using iodoacetamide, a thiol-specific compound that reacts with sulfhydryl groups that are sufficiently nucleophilic, followed by a CHS activity assay at the usual reaction pH. At pH values above the p*K*_a_, the catalytic cysteine is deprotonated and able to react with iodoacetamide, thus inactivating CHS. At pH values below the p*K*_a_, the catalytic cysteine is protonated and protected from iodoacetamide modification, thus retaining CHS activity in the subsequent enzyme assay. The amount of CHS activity remaining after iodoacetamide treatment was expressed as a ratio compared to the CHS activity of a control treatment at the same pH but without iodoacetamide. The p*K*_a_ was calculated using nonlinear regression to fit a log(inhibitor) vs. response equation, which gave the pH at which 50% of maximal inhibition was obtained.

The p*K*_a_ for AtCHS was measured to be 5.428, which is close to the 5.5 measured for MsCHS (Figure 3A). The p*K*_a_ for SmCHS was measured to be 6.468, approximately 1 pH unit higher than that of the two angiosperm CHS orthologs. This elevated p*K*_a_ measured for SmCHS is consistent with the observation of a catalytic cysteine that is less reactive and less prone to oxidation. Also consistent with the crystallographic and plant complementation results, p*K*_a_ values around 5.5 were measured for euphyllophyte orthologs PsCHS and EaCHS, and around 6.5 for the basal-plant orthologs PpCHS (Supplementary Figure 4).

**Figure 3.**
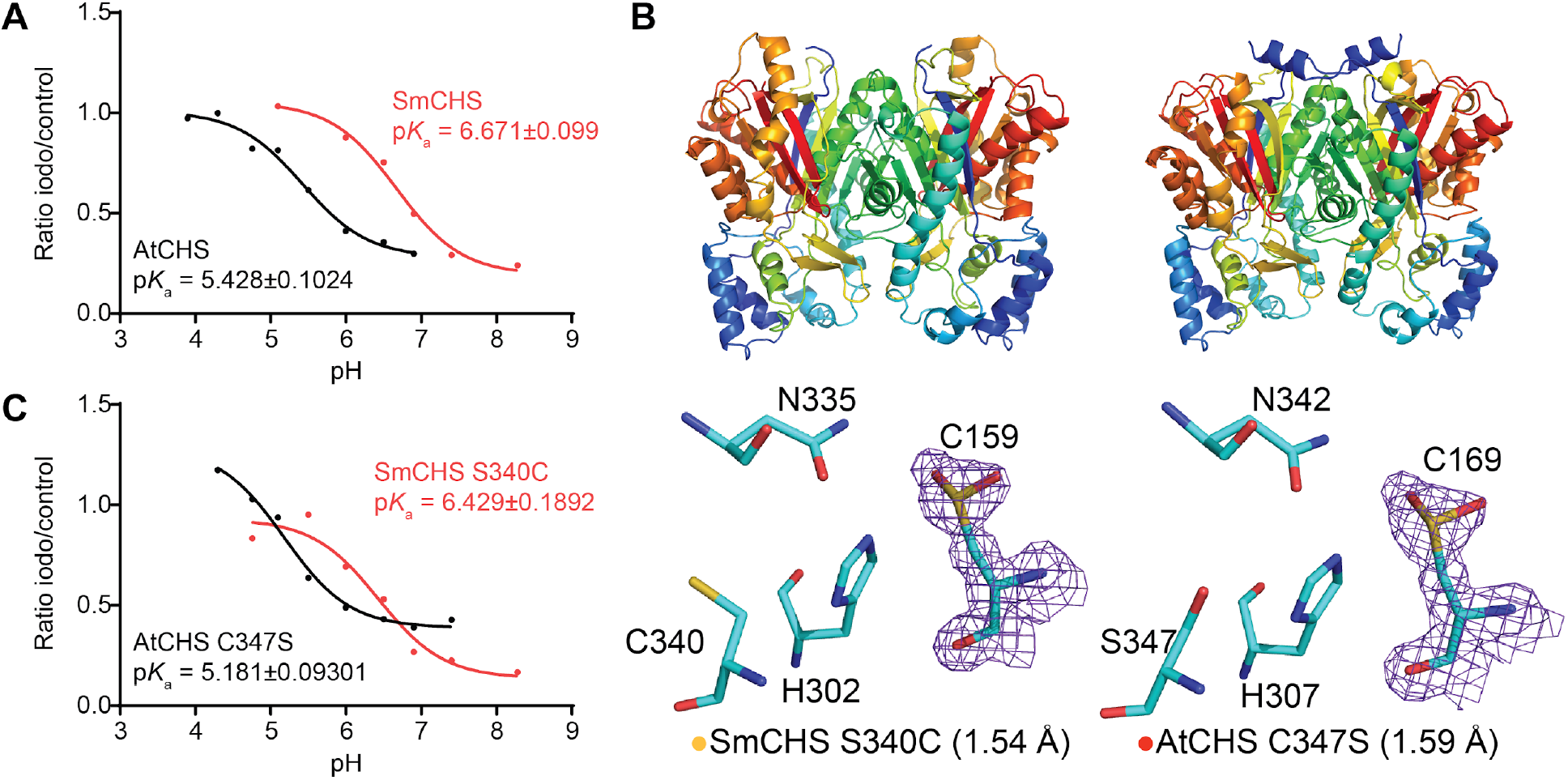
p*K*_a_ measurement of the catalytic cysteine and characterization of key residues that affect p*K*_a_. **A**, p*K*_a_ measurement of AtCHS and SmCHS wild-type enzymes. CHS enzyme was pre-incubated at various pH with or without the 25 µM iodoacetamide inhibitor for 30 s, and an aliquot was taken to run in a CHS activity assay. The ratio of naringenin product produced in the iodoacetamide treatment divided by the control treatment was calculated for each pH point. A nonlinear regression was performed to fit a log(inhibitor) vs. response curve to determine the pH at which 50% of maximal inhibition was achieved, which was determined to be the p*K*_a_ of the catalytic cysteine residue. The p*K*_a_ of AtCHS is close to the 5.5 determined for other euphyllophyte CHSs, whereas the p*K*_a_ of SmCHS is over 1 pH unit higher. **B**, Overall structures and active site configurations of AtCHS C347S and SmCHS S340C single mutants. The 2F_o_-F_c_ electron density map contoured at 1.5σ is shown around the catalytic cysteine. SmCHS S340C shows oxidation of C159, unlike the SmCHS WT. AtCHS C347S has an oxidized C169, like AtCHS WT. **C**, p*K*_a_ measurements of AtCHS C347S and SmCHS S340C mutants.

### Residues near the active-site cavity affect the p*K*_a_ and reactivity of the catalytic cysteine

We next examined the sequence and structural differences between basal-plant and euphyllophyte CHSs that could play a role in modulating catalytic cysteine reactivity. This led us to first identifying a residue near the active site that is conserved as C347 (AtCHS numbering) in AtCHS and other euphyllophyte sequences, and as S340 (SmCHS numbering) in SmCHS and other lycophyte and bryophyte sequences (Figure 2A).

To investigate the role of this residue in modulating catalytic cysteine reactivity, we generated the reciprocal mutations in SmCHS and AtCHS respectively and first characterized these mutant proteins using X-ray crystallography (Figure 3B). Under identical crystallization conditions as wild-type SmCHS, the SmCHS S340C mutant exhibits a partially oxidized catalytic cysteine in its crystal structure, suggesting that the residue does play some role in determining cysteine reactivity. The AtCHS C347S mutant, however, still retains an oxidized catalytic cysteine in its crystal structure.

We then measured the p*K*_a_ of the catalytic cysteine in both SmCHS S340C and AtCHS C347S mutants (Figure 3C). The p*K*_a_ for SmCHS S340C decreases by about 0.25 pH units compared to wild-type SmCHS, consistent with the observation that the SmCHS S340C crystal structure contained a partially oxidized catalytic cysteine. The p*K*_a_ for AtCHS C347S decreases by about 0.25 pH units compared to wild-type AtCHS, also consistent with the observation that the AtCHS C347S crystal structure retained an oxidized catalytic cysteine. Taken together, the crystallographic and p*K*_a_ measurement results suggest that the reciprocal mutation at this residue is not sufficient to act as a simple switch between the active-site environments of euphyllophyte and basal-plant CHSs to modulate catalytic cysteine reactivity. Additional sequence and structural features likely contribute to an active-site environment that lowers the p*K*_a_ of the catalytic cysteine in AtCHS.

To identify these features, we examined a multiple sequence alignment of CHS orthologs from diverse plant species and identified residues that show conserved variations between euphyllophytes and basal-plant lineages (Figure 4A and Supplementary Figure 5). Two residues, F170 and G173 in euphyllophyte CHSs, were found to be substituted as serine and alanine, respectively, in basal-plant lineages. Because of their positions in the alpha helix immediately C-terminal to the catalytic C169, we postulated that these two residues could play a role in determining the structure of the helix, which would have an effect on the electronic environment of the active site, due to the helix dipole’s contribution to lowering the catalytic cysteine p*K*_a_ (Ferrer et al., 1999). Four additional residues near the active-site opening of CHS were also identified as differentially conserved between euphyllophytes and basal plants. We postulated that these positions might affect the dynamics of the active-site tunnel and solvent access to the active site. The six aforementioned residues were mutated in the SmCHS S340C background to their corresponding residues in AtCHS to generate the SmCHS I54M S160F A163G G203S A207Q V258T S340C septuple mutant, termed SmCHS M7. Likewise, the reciprocal mutations were also made in the AtCHS C347S background to generate AtCHS M7.

**Figure 4.**
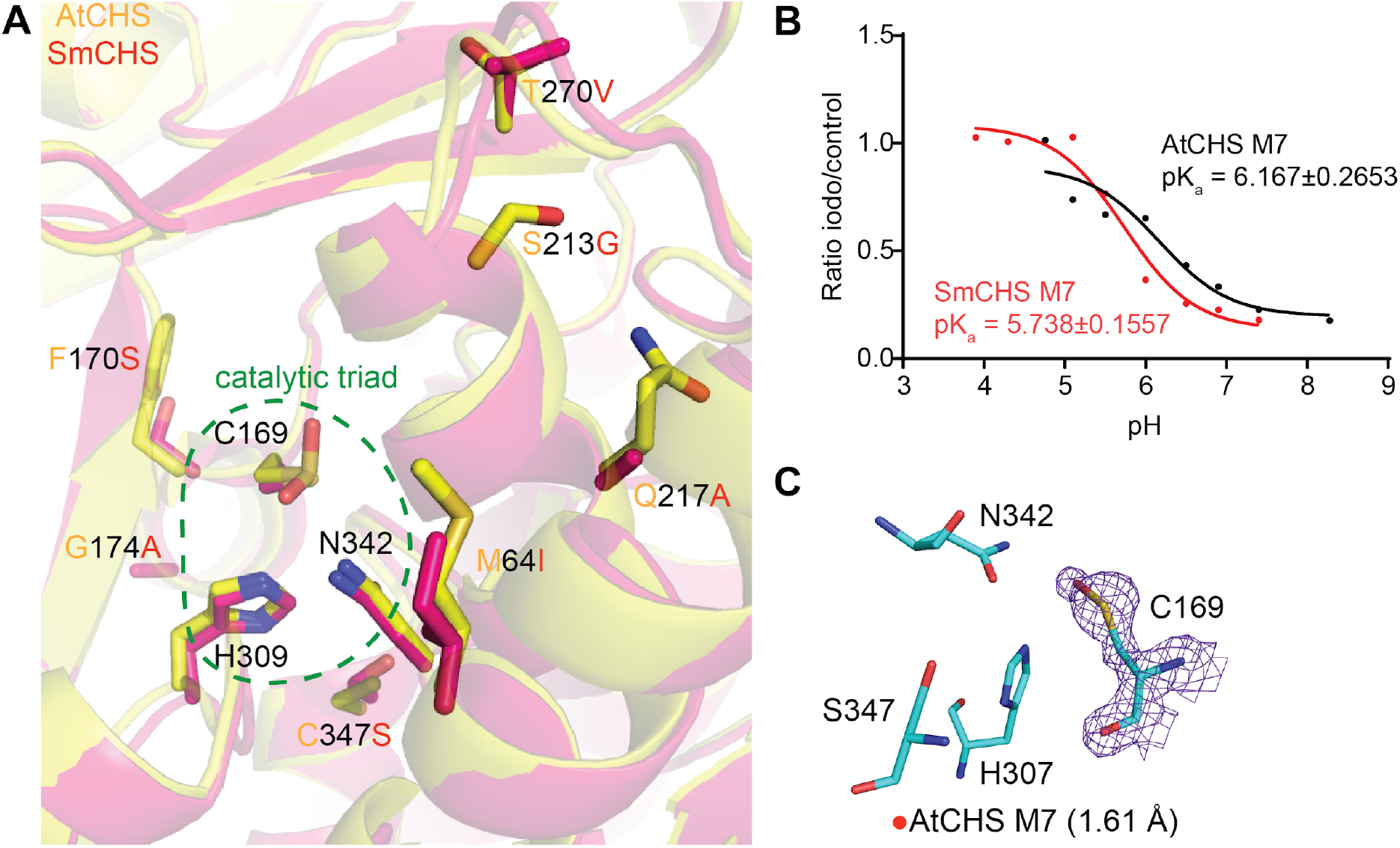
Identification and characterization of additional key residues that affect CHS cysteine reactivity. **A**, Overlaid crystal structures of AtCHS and SmCHS showing the seven conserved residue differences between euphyllophyte and basal-plant CHSs. **B**, p*K*_a_ measurement of AtCHS M7 and SmCHS M7 mutants. The p*K*_a_ of each M7 mutant is about 0.5 pH units higher or lower, respectively, than the corresponding wild type CHS. **C**, The active sites of the two monomers of the AtCHS M7 septuple mutant structure. The 2F_o_-F_c_ electron density map contoured at 1.5σ is shown around the catalytic cysteine. SmCHS S340C shows oxidation of C159, unlike the SmCHS WT. AtCHS C347S has an oxidized C169, like AtCHS WT. **C**, p*K*_a_ measurements of AtCHS C347S and SmCHS S340C mutants. is shown around the catalytic cysteine. The crystal structure shows oxidation to sulfenic acid in the catalytic cysteine of one chain (left) and a reduced cysteine in the other (right).

Compared to SmCHS S340C, the six additional mutations in SmCHS M7 lower the p*K*_a_ by nearly 0.9 pH units from 6.708 to 5.837 (Figure 4). Similarly, the six mutations of AtCHS M7 raise the p*K*_a_ by almost 0.9 pH units from 5.113 to 5.996 compared to AtCHS C347S. Consistent with the p*K*_a_ observation, the dimeric crystal structure of AtCHS M7 has one monomer with a catalytic cysteine singly oxidized to sulfenic acid and one monomer with a reduced cysteine (Figure 4C). Sulfenic acid is more reduced than the doubly oxidized sulfinic acid seen in other euphyllophyte crystal structures, indicating that these six mutations decreased the reactivity of the catalytic cysteine. These mutations represent a part of a possible evolutionary path from ancestral basal-plant CHSs toward the stronger p*K*_a_-lowering properties of euphyllophyte CHSs. Any further attempts at engineering CHS to fully swap the p*K*_a_-lowering properties between AtCHS and SmCHS would likely require different methods of searching for conserved sequence differences, beyond visual observation of structural differences. An analysis of the CHS multiple sequence alignment using ancestral sequence reconstruction with FastML (Ashkenazy et al., 2012) identified eight additional positions that are differently conserved between euphyllophytes and basal plants and could affect CHS function based on their position in the CHS crystal structure (Supplementary Figure 6).

### Molecular dynamics simulations reveal differences in active-site interactions between basal-plant and euphyllophyte CHSs

Our crystal structures revealed a correlation between the p*K*_a_ of the catalytic cysteine and a set of residues near the active site. To further investigate the mechanisms underlying these conserved differences between euphyllophyte and basal-plant CHSs, we employed molecular dynamics (MD) simulations to examine the interactions between these residues. We first surveyed the potential role of the C347S substitution (AtCHS numbering) in affecting the active site environment in wild-type AtCHS and SmCHS (Figure 5A). In wild-type AtCHS, where the largest cluster represents 70.3% of all structures sampled in this simulation, the thiol group of C347 points away from the active site and cannot form any stable interaction with the catalytic H309 (distance 6.5 Å). In contrast, the corresponding S340 in SmCHS is 2.8 Å away from the histidine in the largest cluster, representing 98.7% of all structures sampled in the SmCHS simulation.

**Figure 5.**
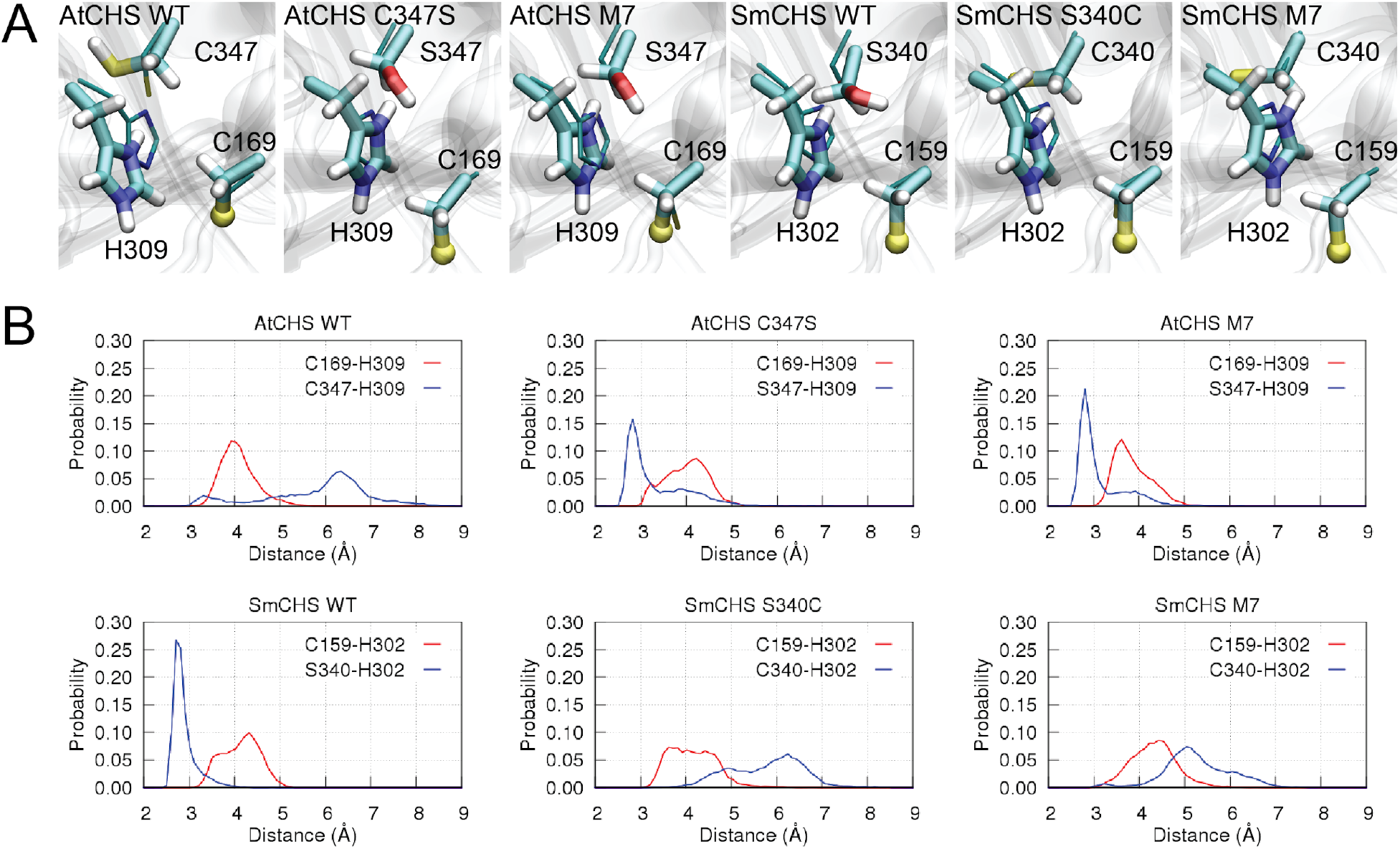
Molecular dynamics simulations of CHS orthologs and mutants. **A**, The centroid structure of the largest cluster of the catalytic pair C169-H309. For visualization purposes, the sulfur atom in the ionic cysteine C169 is shown in a ball representation, and crystal structures are depicted as thin sticks. **B**, Distributions of inter-residue distances obtained from simulations.

Next, we determined the inter-residue distances between the ionic pair C169-H309 as in AtCHS or C159-H302 as in SmCHS and between residue C347 (AtCHS)/S340 (SmCHS) and the catalytic histidine (Figure 5B). For wild-type SmCHS simulation, we observe a sharp peak at around 2.8 Å between S340 and H302, reflecting a stable hydrogen bond between the two residues. On the contrary, no such short-distance peak is observed for wild- type AtCHS. These results suggest that the catalytic histidine is stabilized upon forming a hydrogen bond with S340 in SmCHS, but such an interaction is relatively loose in AtCHS. Similar differences between the other euphyllophyte and basal-plant CHSs are also seen for PsCHS, EaCHS, and PpCHS (Supplementary Figure 7).

To further investigate the motion of the catalytic histidine in various mutant enzyme active-site environments, we also performed MD simulations of AtCHS C347S, SmCHS S340C, AtCHS M7, and SmCHS M7 (Figure 5A). The largest cluster sizes were 86.0%, 66.6%, and 96.7%, and 71.0%, respectively. S347 and H309 in AtCHS mutants adopt a similar conformation to wild-type SmCHS. In contrast, no stable hydrogen bond between C340 and H302 is formed in the largest cluster of the SmCHS mutant simulations. Introducing point mutations dramatically changes the distributions of those key inter-residue distances. In the AtCHS C347S mutant, the S347-H309 distance dramatically shortens to a peak around 2.8 Å, and introducing the six additional mutations in AtCHS M7 further increases the height of the peak. This suggests that mutating these seven positions in AtCHS to the corresponding residues in SmCHS can allow the active-site residues to approximate the interactions of wild-type SmCHS. The opposite effect is seen in SmCHS S340C and SmCHS M7 mutants, which recapitulate the weak interaction between C347 and H309 seen in wild-type AtCHS.

Based on these results, we hypothesize that the strong S340-H302 interaction facilitated by the SmCHS active site environment may weaken the stabilizing effect of H302 on the catalytic cysteine thiolate compared to that in AtCHS, thus contributing to the higher p*K*_a_. Meanwhile, the inter-residue distance of the catalytic cysteine-histidine ionic pair is rather stable in all CHS simulations, ranging from 3 to 5 Å and centered at around 4.1 Å. This suggests that the C347S substitution (AtCHS numbering) does not directly break this ionic interaction but may subtly influence the charge distribution on the histidine imidazole ring to perturb the catalytic cysteine p*K*_a_ (Figure 6). In addition, the presence of a cysteine appears to decrease solvent content in the active site compared to serine, which would increase the p*K*_a_-lowering effect of the ionic interaction between histidine and the catalytic cysteine (Supplementary Note and Supplementary Figure 8). Taken together, our results suggest that euphyllophyte CHSs have evolved to enhance the reactivity of the catalytic cysteine through the modification of specific interactions between active-site residues to allow for stronger stabilization of the thiolate.

**Figure 6.**
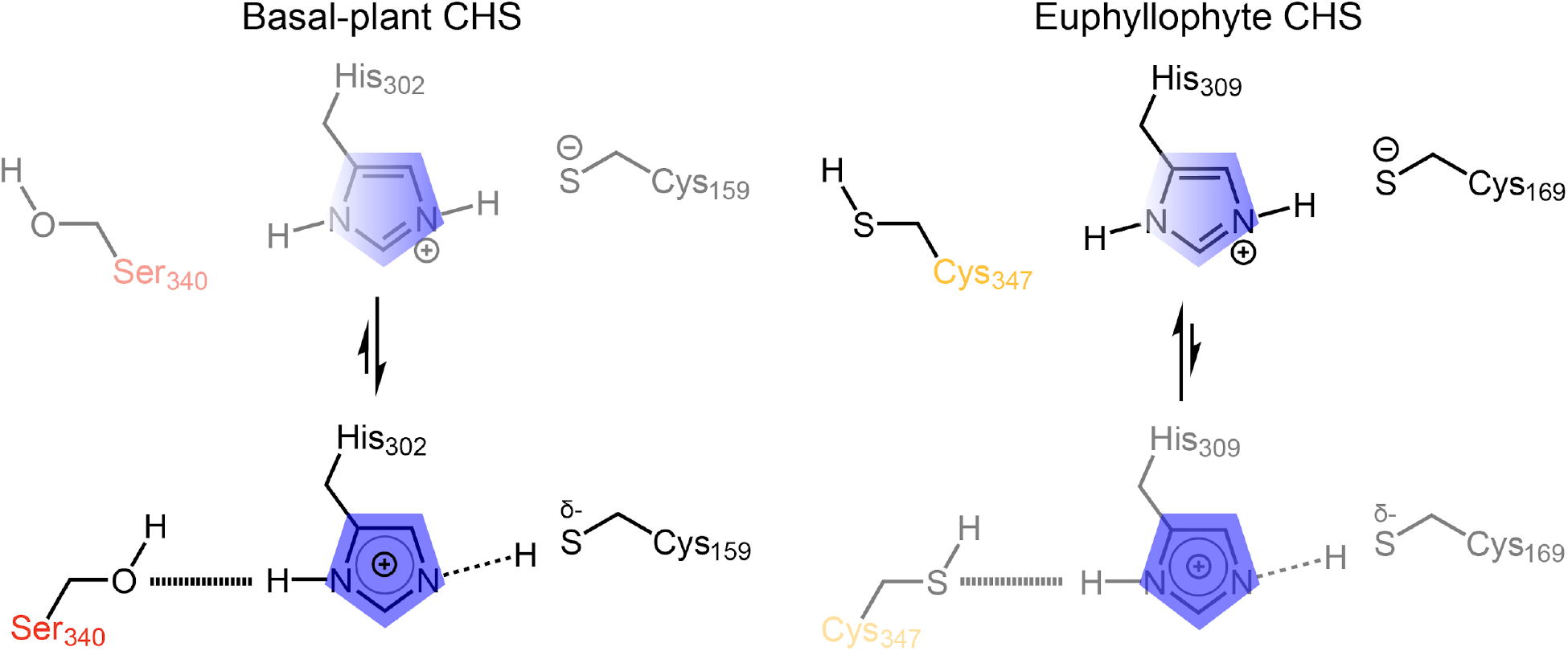
Proposed model for differential modulation of catalytic cysteine nucleophilicity in basal-plant (left) and euphyllophyte (right) CHSs. In basal-plant CHSs (left), the serine (S340 in SmCHS) interacts more strongly with the histidine of the catalytic triad, weakening the ionic interaction that stabilizes the thiolate form of the catalytic cysteine. This is depicted as a shift of the equilibrium toward a state in which the positive charge on the histidine (blue) is shifted away from the catalytic cysteine (C159 in SmCHS) and the shared proton interacts more closely with cysteine. In euphyllophyte CHSs (right), this position mutated to a cysteine (C347 in AtCHS), which interacts relatively loosely with the catalytic histidine, in turn strengthening the ionic interaction between the catalytic histidine and the activated thiolate of the catalytic cysteine. This is depicted as a shift of the equilibrium toward a state in which the positive charge on the histidine (blue) is shifted toward the catalytic cysteine (C169 in AtCHS).

## Discussion

As early plants initially migrated from water to land and further radiated to occupy diverse terrestrial environmental niches, they continuously encountered new challenges from biotic and abiotic stresses. The greatly expanded diversity and increased abundance of flavonoids in certain plant lineages could have increased the demand for metabolic flux into flavonoid biosynthesis. One adaptive strategy to meet this demand, among many others, is to increase the enzymatic efficiency of chalcone synthase, the first committed enzyme of flavonoid biosynthesis that gates flux from general phenylpropanoid metabolism. One property of CHS that affects its enzymatic efficiency is the reactivity of the first step of nucleophilic attack on *p*-coumaroyl-CoA. To investigate this, we performed structural, biochemical, mutagenesis, and molecular dynamics experiments on CHS orthologs from five major plant lineages. Our results suggest that euphyllophyte CHSs have indeed evolved new structural features to increase the reactivity of their catalytic cysteine compared to basal-plant CHSs.

To identify sequence and structural features between euphyllophyte and basal-plant CHSs that lead to this difference in enzymatic properties, we generated mutants in the background of AtCHS and SmCHS at various positions with conserved sequence differences segregating euphyllophyte and basal-plant CHSs. AtCHS M7 and SmCHS M7 had p*K*_a_ values raised by about 0.7 and lowered by about 0.9 pH units from the wild-type enzymes, respectively. Furthermore, AtCHS M7 also exhibits a less oxidized catalytic cysteine in its crystal structure than in wild-type AtCHS. These results indicate that we were able to identify residue changes that partially traced the evolutionary path from SmCHS to AtCHS that increased the reactivity of the catalytic cysteine. In the type III PKS family, the introduction of a large number of mutations to yield subtle changes in enzyme activity is not unprecedented. Stilbene synthase (STS) produces resveratrol, a tetraketide product whose biosynthetic mechanism differs from that of naringenin chalcone in only the final cyclization step. In a previous study, a total of 18 point mutations were required to convert CHS activity to STS activity, through small changes in the hydrogen-bonding network in the active site (Austin et al., 2004).

To examine in detail the intramolecular interactions that lead to enhanced cysteine reactivity, we performed molecular dynamics simulations on CHS. In comparing different CHS orthologs and point mutants, we observed that the presence of a cysteine in the nearby position leads to a weak interaction between the cysteine and histidine, as indicated by the broad distribution of inter-residue distances centered at a distance greater than 5 Å, too long for a stable hydrogen bond. In contrast, when a serine is present, the sharp peak of serine-histidine inter-residue distance around 2.75 Å suggests the presence of a strong hydrogen bond. This hydrogen bonding likely shifts the electron density of the histidine away from the catalytic cysteine, weakening the imidazoline-thiolate ion pair. This weakened ionic interaction would lead to less p*K*_a_ depression compared to CHS orthologs and mutants containing a cysteine in the nearby position, where the histidine is able to maintain a stronger ion pair with the catalytic cysteine and lower the p*K*_a_ to a greater degree. This is reminiscent of the role of aspartate 158 in papain, a cysteine protease that also uses a cysteine-histidine-asparagine catalytic triad for nucleophilic attack on its substrates (Storer and Ménard, 1994). Although D158 is not essential for papain activity, its side chain affects the pH-activity profile by forming a hydrogen bond with the backbone amide of the catalytic histidine. This interaction stabilizes the catalytic ionic pair and maintains an optimal orientation of active-site residues. A D158E mutant papain had a pH-activity profile shifted by 0.3 pH units, about the same magnitude of the effect we observed on p*K*_a_ for CHS cysteine/serine mutants.

We propose a model of the role of position 347 in enhancing CHS reactivity (Figure 6). In the basal example of SmCHS, the serine interacts more strongly with the histidine of the catalytic triad, weakening the ionic interaction that stabilizes the thiolate form of the catalytic cysteine. In euphyllophyte CHSs, this position mutated to a cysteine, which interacts more poorly with the histidine, strengthening the ionic interaction and stabilizing the activated thiolate of the catalytic cysteine.

While the mechanism of how the other six mutations in the M7 mutants affect the catalytic cysteine is not entirely clear, we noticed that, possibly due to the smaller side chains of the S213G and Q217A mutations, AtCHS M7 has a surface helix in a slightly different conformation than wild-type AtCHS, leading to a slightly wider active-site opening. There is also a newly solvent-accessible cavity as determined by a computational cavity-finding software (Supplementary Figure 8). These structural differences could lead to subtle changes in the amino acid backbone dynamics near the active site and thus alter the active-site volume or electronic environment, which could alter the p*K*_a_ of the catalytic cysteine (Jez and Noel, 2000).

Although cysteine sulfenic and sulfinic acid have been thought of as crystallographic artifacts, an increasing number of studies have shown that this type of cysteine oxidation can play an important functional role. In particular, cysteine sulfinic acid has been shown to play a regulatory role in reversible inhibition of the activity of enzymes such as protein tyrosine phosphatase 1B and glyceraldehyde-3-phosphate dehydrogenase, suggesting that cysteines sensitive to oxidation can be an evolved trait (van Montfort et al., 2003; Peralta et al., 2015).

Our results demonstrate that euphyllophytes could have evolved a CHS enzyme that is intrinsically more active, with increase cysteine reactivity as one component, as one adaptation to produce the larger suite of flavonoids needed to counter the various environmental stresses they face. Although it may seem counterintuitive for euphyllophytes, which encounter more oxidative environments than do basal plants, to rely on a CHS enzyme that is more susceptible to oxidation, this susceptibility may be an unavoidable trade-off resulting from the chemical nature of a more nucleophilic cysteine: a catalytic cysteine more reactive toward substrate is also more reactive toward oxidants like hydrogen peroxide. To compensate for this increased susceptibility to oxidation, euphyllophytes may have evolved other systems to better maintain the redox environment inside the cell, one of those systems being the antioxidant flavonoids themselves.

## Materials And Methods

### Cloning and site-directed mutagenesis of CHSs

Total RNA was obtained from *Arabidopsis thaliana, Pinus sylvestris, Equisetum arvense, Selaginella moellendorffii*, and *Physcomitrella patens*. Reverse transcription was performed to obtain cDNA. The open reading frames (ORFs) of five *CHS* orthologs were amplified via polymerase chain reaction (PCR) from cDNA, digested with NcoI and XhoI, and ligated into NcoI- and XhoI-digested pHis8–3 or pHis8–4B *Escherichia coli* expression vectors. Site-directed mutagenesis was performed according to the QuikChange II Site-Directed Mutagenesis protocol (Agilent Technologies).

### Transgenic Arabidopsis

The At*CHS* promoter (defined as 1328 bp of sequence upstream of the CHS transcription start site) was amplified via PCR from Arabidopsis genomic DNA, digested with HindIII and XhoI, and ligated into HindIII- and XhoI-digested pCC 1136, a promoterless Gateway cloning binary vector containing a BAR resistance gene marker, to generate pJKW 0152. The five *CHS* ORFs described above were then PCR amplified from cDNA and cloned into pCC 1155, an ampicillin-resistant version of the pDONR221 Gateway cloning vector, with BP clonase in the Gateway cloning method (Thermo-Fisher). The resulting vectors were recombined with pJKW 0152 using LR clonase in the Gateway cloning method to generate the final binary constructs. *Agrobacterium tumefaciens*-mediated transformation of Arabidopsis was performed using the floral dipping method (Weigel and Glazebrook 2002).

### Recombinant protein expression and purification

CHS genes were cloned into pHis8–3 or pHis8–4B, bacterial expression vectors containing an N-terminal 8×His tag followed by a thrombin or tobacco etch virus (TEV) cleavage site, respectively, for recombinant protein production in *E. coli*. Proteins were expressed in the BL21(DE3) *E. coli* strain cultivated in terrific broth (TB) and induced with 0.1 mM isopropyl β-D-1-thiogalactopyranoside (IPTG) overnight at 18 °C. *E. coli* cells were harvested by centrifugation, resuspended in 150 mL lysis buffer (50 mM Tris pH 8.0, 500 mM NaCl, 30 mM imidazole, 5 mM DTT), and lysed with five passes through an M-110L microfluidizer (Microfluidics). The resulting crude protein lysate was clarified by centrifugation (19,000 g, 1 h) prior to QIAGEN nickel–nitrilotriacetic acid (Ni–NTA) gravity flow chromatographic purification. After loading the clarified lysate, the Ni–NTA resin was washed with 20 column volumes of lysis buffer and eluted with 1 column volume of elution buffer (50 mM Tris pH 8.0, 500 mM NaCl, 300 mM imidazole, 5 mM DTT). 1 mg of His-tagged thrombin or TEV protease was added to the eluted protein, followed by dialysis at 4 °C for 16 h in dialysis buffer (50 mM Tris pH 8.0, 500 mM NaCl, 5 mM DTT). After dialysis, the protein solution was passed through Ni–NTA resin to remove uncleaved protein and His-tagged TEV. The recombinant proteins were further purified by gel filtration on an ÄKTA Pure fast protein liquid chromatography (FPLC) system (GE Healthcare Life Sciences). The principal peaks were collected, verified by SDS–PAGE, and dialyzed into a storage buffer (12.5 mM Tris pH 8.0, 50 mM NaCl, 5 mM DTT). Finally, proteins were concentrated to > 10 mg/mL using Amicon Ultra-15 Centrifugal Filters (Millipore).

### Protein crystallization

All protein crystals were grown by hanging drop vapor diffusion at 4 °C, except for EaCHS at 20 °C. For AtCHS wild-type and C347S crystals, 1 µL of 10 mg/mL protein was mixed with 1 µL of reservoir solution containing 0.1 M HEPES (pH 7.5), 0.3 M ammonium acetate, 14% (v/v) PEG 8000, and 5 mM DTT. For AtCHS M7 crystals, 1 µL of 16.33 mg/mL protein was mixed with 1 µL of reservoir solution containing 0.125 NaSCN, 20% (v/v) PEG 3350, and 5 mM DTT; 0.2 µL of a crystal seed stock from previous rounds of crystal optimization was also added. For SmCHS wild-type and S340C crystals, 1 µL of 10 mg/mL protein was mixed with 1 µL of reservoir solution containing 0.1 M MOPSO (pH 6.6), 0.3 M Mg(NO_3_)_2_, 19% (v/v) PEG 4000, and 5 mM DTT. For EaCHS, 1 µL of 16.92 mg/mL protein was mixed with 1 µL of reservoir solution containing 0.15 M LiCl, 8% PEG 6000, and 5 mM DTT. For PsCHS, 1.66 µL of 14.65 mg/mL protein was mixed with 0.67 µL of reservoir solution containing 0.14 M NH_4_Cl, 20% (v/v) PEG 3350, and 5 mM DTT; 0.2 µL of a crystal seed stock from previous rounds of crystal optimization was also added. For PpCHS, 1 µL of 10 mg/mL protein was mixed with 1 µL of reservoir solution containing 0.1 M MES (pH 6.9), 18% (v/v) PEG 20000, and 5 mM DTT. Crystals were harvested within 1 week and transferred to a cryoprotection solution of 17% glycerol and 83% reservoir solution. H_2_O_2_ soaking of SmCHS crystals was performed by adding H_2_O_2_ to 1 mM to the cryoprotection solution and incubating at 4 °C for 75 min. Single crystals were mounted in a cryoloop and flash-frozen in liquid nitrogen.

### X-ray diffraction and structure determination

X-ray diffraction data were collected at beamlines 8.2.1 and 8.2.2 of the Advanced Light Source at Lawrence Berkeley National Laboratory on ADSC Quantum 315 CCD detectors for AtCHS wild-type, AtCHS C347S, and SmCHS S340C crystals. X-ray diffraction data were collected at beamlines 24-ID-C and 24-ID-E of the Advanced Photon Source at Argonne National Laboratory on an ADSC Quantum 315 CCD detector, Eiger 16M detector, or Pilatus 6M detector for SmCHS wild-type, EaCHS, PsCHS, and AtCHS M7 crystals. Diffraction intensities were indexed and integrated with iMosflm (Battye et al., 2011) and scaled with Scala under CCP4 (Evans, 2006; Winn et al., 2011). The phases were determined with molecular replacement using Phaser under Phenix (Adams et al., 2010). Further structural refinement utilized Phenix programs. Coot was used for manual map inspection and model rebuilding (Emsley and Cowtan, 2004). Crystallographic calculations were performed using Phenix.

### Comparative sequence and structure analyses

CHS protein sequences were derived from NCBI and the 1000 Plants (1KP) Project (Matasci et al., 2014; NCBI Resource Coordinators, 2016). In all cases, AtCHS was used as the search query. Amino acid alignment of CHS orthologs was created using MUSCLE with default settings (Edgar, 2004). UCSF Chimera and ESPript were used to display the multiple-sequence alignments shown in Figure 2, Supplementary Figure 5, and Supplementary Figure 6 (Pettersen et al., 2004; Robert and Gouet, 2014). Phylogenetic analysis was performed using MEGA7 (Kumar et al., 2016). All structural figures were created with the PyMOL Molecular Graphics System, version 1.3 (Schrödinger, LLC) (DeLano, 2016). Active site cavity measurements for the AtCHS and AtCHS M7 structures were determined using KVFinder (Oliveira et al., 2014).

### Enzyme assays and p*K*_a_ measurement

A 4CL-CHS coupled assay was used for kinetic analysis. A 4CL reaction master mix was made by incubating 917 nM Arabidopsis thaliana 4CL1 (NCBI accession number NP_175579.1) in 100 mM Tris-HCl (pH 8.0), 5 mM MgCl_2_, 5 mM ATP, 100 µM *p*-coumaric acid, 100 µM coenzyme A, and 10 or 50 µM malonyl-CoA for 30 min at room temperature to generate *p*-coumaroyl-CoA at a final concentration of 70 µM. This 4CL was divided into individual aliquots of 196 µL in Eppendorf tubes. CHS enzyme was incubated for 30 or 60 s in 16 µL volumes using a triple buffer system (50 mM AMPSO, 50 mM sodium phosphate, 50 mM sodium pyrophosphate, various pH) (Ellis and Morrison, 1982) (Schlegel et al., 1998) at room temperature in the presence of 25 µM iodoacetamide for the inactivation sample or water for the control sample. Aliquots (4 µL) were withdrawn from the incubation mixture and added to the standard coupled CHS assay system. The CHS reaction was run for 10 min at room temperature and stopped by addition of 200 µL methanol.

The assay samples were centrifuged and analyzed directly by liquid chromatography−mass spectrometry (LC−MS). LC was conducted on a Dionex UltiMate 3000 UHPLC system (Thermo Fisher Scientific), using water with 0.1% formic acid as solvent A and acetonitrile with 0.1% formic acid as solvent B. Reverse phase separation of analytes was performed on a Kinetex C18 column, 150 × 3 mm, 2.6 µm particle size (Phenomenex). The column oven was held at 30 °C. Samples were eluted with a gradient of 5–60% B for 9 min, 95% B for 3 min, and 5% B for 3 min, with a flow rate of 0.7 mL/min. MS analysis was performed on a TSQ Quantum Access Max mass spectrometer (Thermo Fisher Scientific) operated in negative ionization mode with a SIM scan centered at 271.78 m/z to detect naringenin chalcone.

The pH profiles (pH on the X-axis, ratio of naringenin chalcone produced with iodoacetamide-treatment to control on the Y-axis) were determined by fitting raw data to the log(inhibitor) vs. response equation using nonlinear regression in Prism, version 6.0f (GraphPad Software).

### Molecular dynamics

All MD simulations were performed using the GROMACS 5.1.4 package (Abraham et al., 2015) and CHARMM force field (Best et al., 2012). The catalytic residues were modeled as protonated histidine (H309 in AtCHS number) and deprotonated cysteine (C169 in AtCHS numbering). All CHSs were constructed as dimers and were pre-aligned to the wild-type AtCHS crystal structure using the Multiseq plugin of VMD (Roberts et al., 2006). All CHS dimers were solvated with 0.1 M NaCl in a dodecahedron box. Before the production runs, all systems were submitted to a minimization, followed by a 500-ps NVT and a 500-ps NPT run with heavy atoms constrained. This was followed by another 5-ns NPT simulation with protein backbone constrained. In all simulations, an integration time step of 2 fs was used, with bonds involving hydrogens constrained using LINCS (Hess, 2008; Hess et al., 1997). The van der Waals interaction was smoothly switched off starting from 10 Å, with a cut-off distance of 12 Å. The neighboring list was updated every 10 steps with Verlet cutoff-scheme. The electrostatic interaction was evaluated using Particle-Mesh-Ewald (PME) summation (Darden et al., 1993) with a grid spacing of 1.5 Å to account for the long-range interaction, while its short-range interaction in real space had a cut-off distance of 12 Å. The velocity-rescaling thermostat (Bussi et al., 2007) and Parrinello-Rahman barostat (Nosé and Klein, 1983; Parrinello and Rahman, 1981) were employed to maintain the temperature at 300 K and the pressure at 1 bar.

For each CHS, three copies of 200-ns production runs were performed. The aggregated simulation time of all CHS wildtype and mutant systems is 5.4 µs. The two monomers of a given CHS were treated equivalently in the analysis; i.e., the three copies of trajectories of each monomer were combined after they were aligned to chain A of the associated crystal structure, resulting in a total of 1.2-µs trajectory for analysis of a given CHS system. Clustering analysis was carried out with GROMACS gmx cluster with a RMSD cutoff of 0.1 nm. The inter-residue distance was measured using the tcl scripting abilities provided by VMD (Humphrey et al., 1996). The minimum distance between the two nitrogen atoms of the catalytic histidine and the associated hydroxyl, thiol, or thiolate group of its serine or cysteine partener was taken as the inter-residue distance. Water occupancy calculation was performed using the volmap plugin of VMD (Humphrey et al., 1996).

### Author Contributions

G.L. and J.K.W. designed and performed all experiments and analyzed data. Y.C.C. and Y.W. performed the molecular dynamics simulations. All authors participated in writing the paper.

## Acknowledgements

We thank Joseph Noel and members of the Noel lab of the Salk Institute for early discussion regarding catalytic cysteine oxidation the Noel lab first observed in various plant type III PKSs, which motivated this work. This work was supported by the Howard Hughes Medical Institute, the National Science Foundation (CHE-1709616, J.K.W.), Pew Scholars Program in the Biomedical Sciences (grant number 27345, J.K.W.), the Searle Scholars Program of the Kinship Foundation (grant number 15-SSP-162, J.K.W.), and direct grants from the Chinese University of Hong Kong (Y.W.). This work is based on research conducted at the Northeastern Collaborative Access Team (NE-CAT) beamlines, which are funded by the National Institute of General Medical Sciences from the National Institutes of Health (P41 GM103403). The Pilatus 6M detector on NE-CAT 24-ID-C beam line is funded by an NIH-ORIP HEI grant (S10 RR029205). This research used resources of the Advanced Photon Source, a U.S. Department of Energy (DOE) Office of Science User Facility operated for the DOE Office of Science by Argonne National Laboratory under Contract No. DE-AC02–06CH11357.

## Supplementary Information

### Supplementary Note

Our MD calculations show that the C347S substitution (AtCHS numbering) can significantly affect active-site solvation. The occupancy of water molecules within the active site was measured with a resolution of 1 Å^3^ (Supplementary Figure 6). Interestingly, S347 in AtCHS C347S and M7 mutants attracts more water toward itself and H309. Similarly, the wild-type SmCHS is also considerably wetter than the wild-type AtCHS: employing a cylinder with a radius of 9 Å and a height of 13 Å to enclose the catalytic residues, we found that the average number of water molecules enclosed was 40.0 for SmCHS and 31.4 for AtCHS. The ability of a serine to attract more water is also observed in simulations of EaCHS, PpCHS, and PsCHS, although in SmCHS mutants the active site remains rather wet despite the mutation of serine to cysteine (Supplementary Figure 6).

AtCHS M7 also showed a wider active-site opening than wild-type AtCHS, which may also affect solvent access to the active site, as shown by the large cavity found in cavity analysis. In addition to changing the hydrogen bonding network, the decreased solvation in euphyllophyte CHSs would enhance the p*K*_a_-lowering effect of the histidine, because ionic effects are enhanced as the dielectric constant decreases along with solvent polarity (Harris and Turner, 2002).

---

**Table S1.**
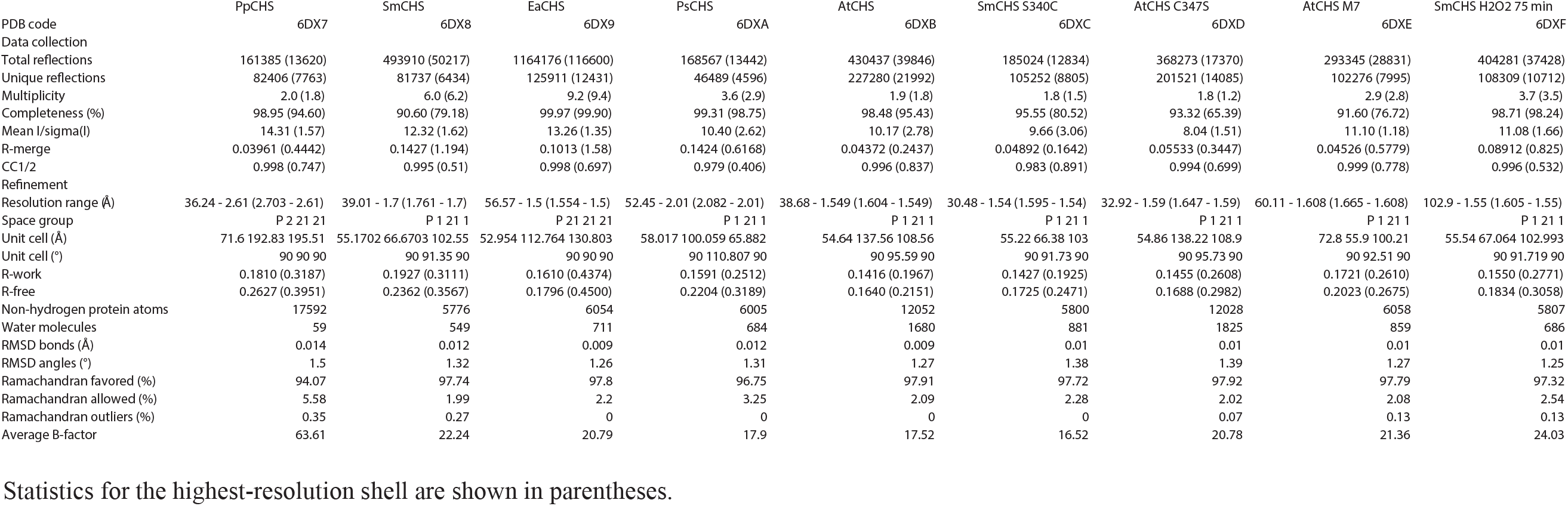
Crystallographic data collection and refinement statistics.

## Supplementary Figures

**Supplementary Figure 1.**
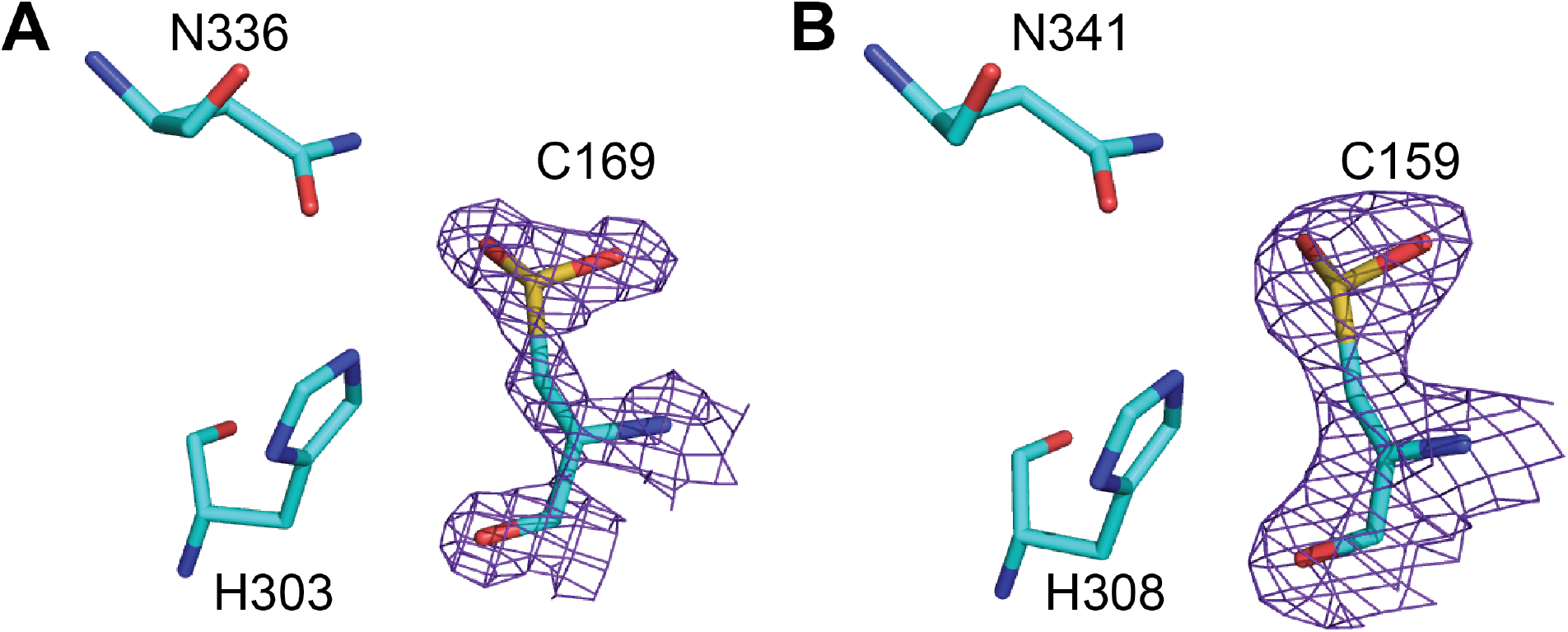
Active site structures of *Medicago sativa* CHS (**A**) and *Gerbera hybrida* 2-pyrone synthase (**B**) showing catalytic cysteine oxidized to sulfinic acid. The 2F_o_-F_c_ composite map contoured at 1.5σ is shown around the catalytic cysteine.

**Supplementary Figure 2.**
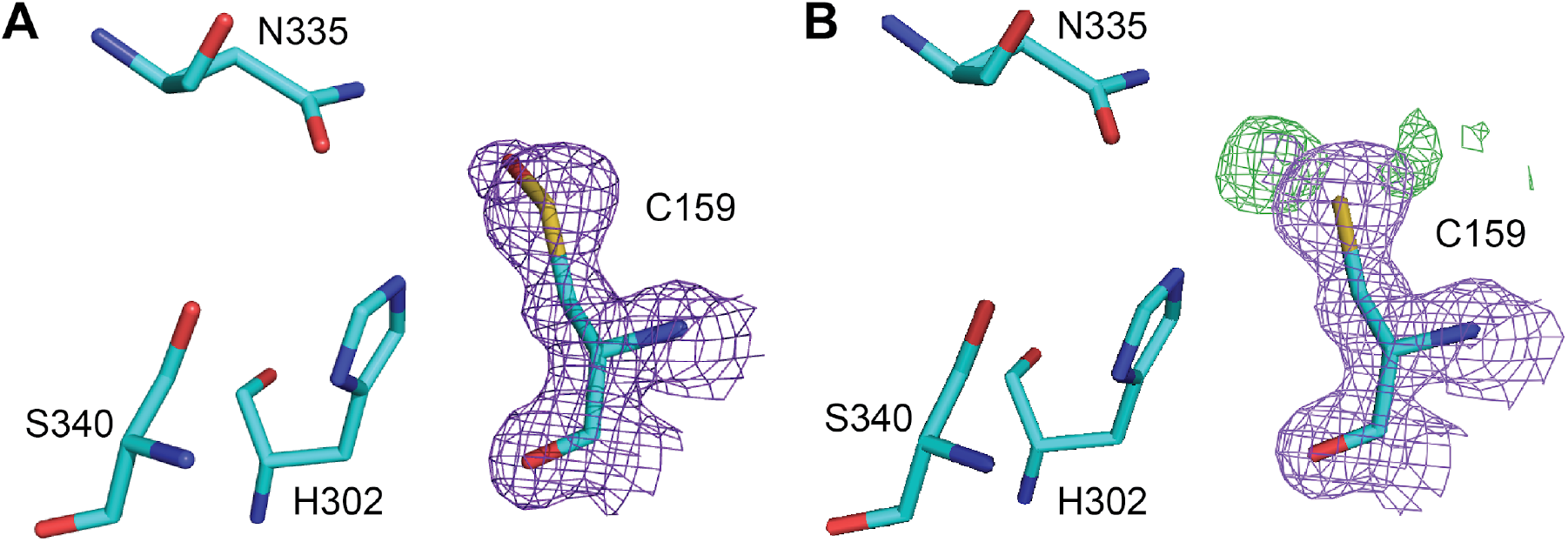
Active site structure of SmCHS crystals soaked in 1 mM hydrogen peroxide for 75 min. **A**, The 2F_o_-F_c_ composite map to 1.55 Å resolution and contoured at 1.5σ is shown around the catalytic cysteine, modeled as oxidized to sulfenic acid. **B**, The 2F_o_-F_c_ composite map to 1.55 Å resolution and contoured at 1.5σ is shown as purple and the F_o_-F_c_ difference map contoured at 3.0σ is shown as green around the catalytic cysteine, modeled as reduced cysteine, indicating clear residual electron density for the oxidized sulfenic acid.

**Supplementary Figure 3.**
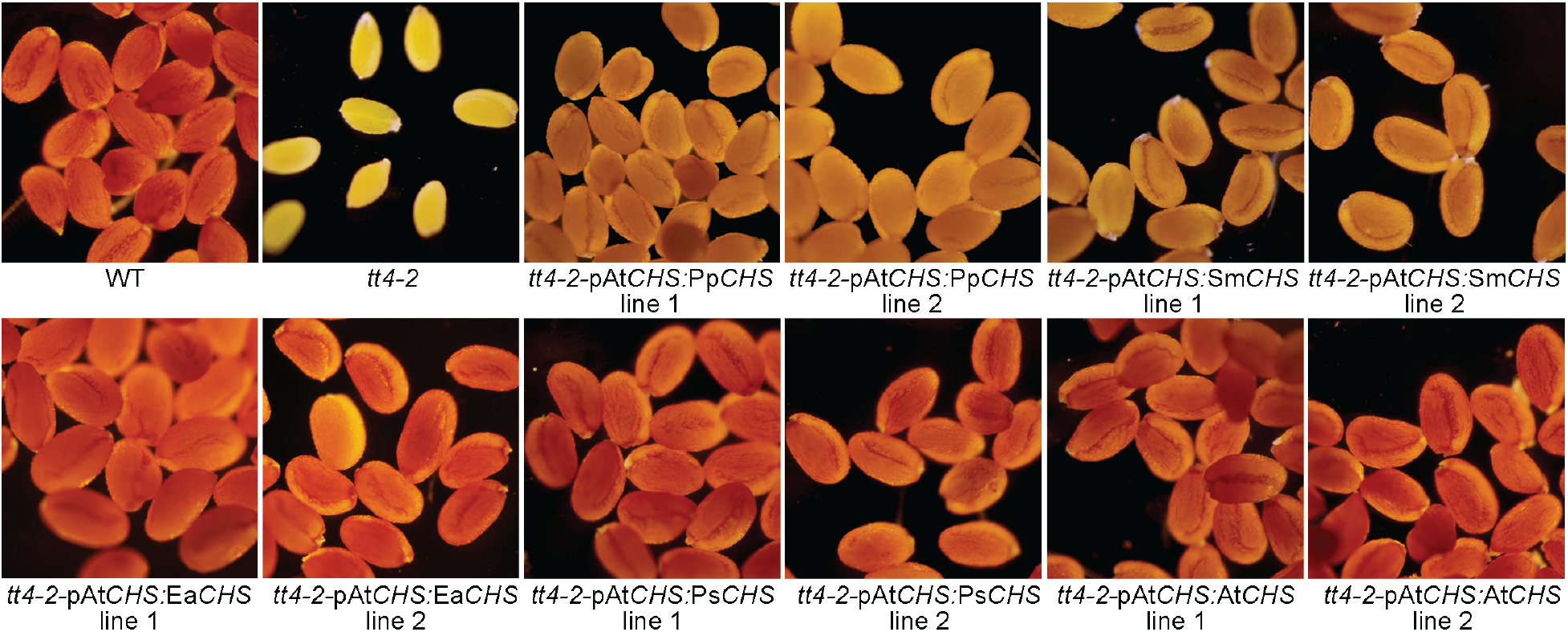
Complementation of the *transparent testa* seed phenotype of *tt4–2* mutant *Arabidopsis thaliana*. CHS orthologs were expressed under the At*CHS* promoter. CHS from euphyllophytes (AtCHS, PsCHS, EaCHS) fully complement the mutant phenotype, whereas CHS from basal land plants (SmCHS, PpCHS) only partially complement.

**Supplementary Figure 4.**
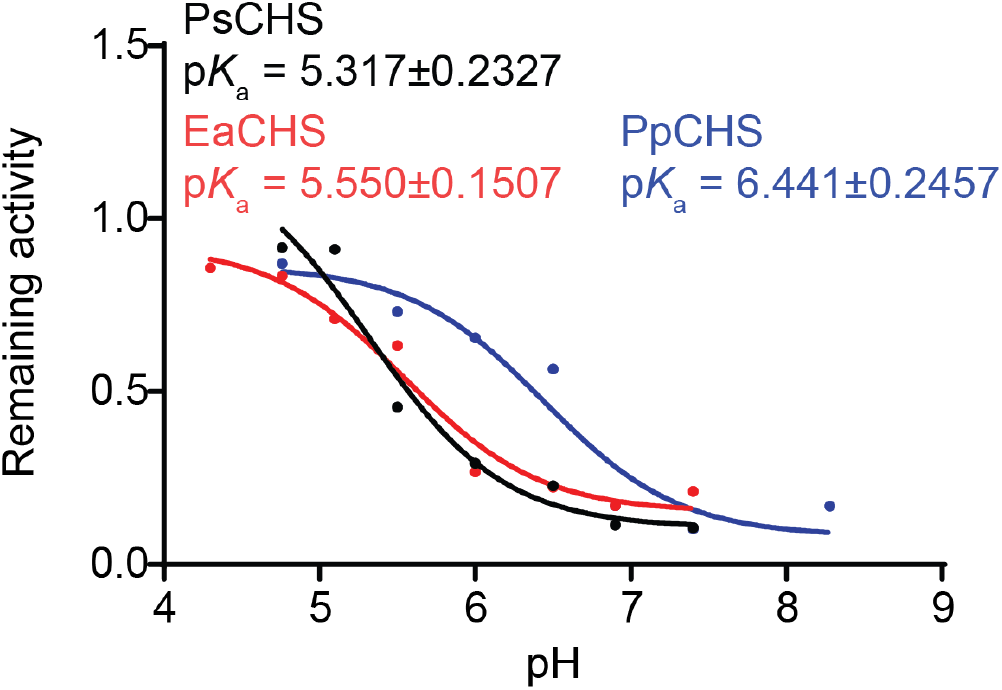
p*K*_a_ measurement of PsCHS, EaCHS, and PpCHS wild type enzymes. CHS enzyme was pre-incubated at various pH in the 25 µM iodoacetamide inhibitor or water (control) for 30 s, and an aliquot was taken to run in a CHS activity assay. The ratio of naringenin product produced in the iodoacetamide treatment divided by the control treatment was calculated for each pH point. A nonlinear regression was performed to fit a log(inhibitor) vs. response curve to determine the pH at which 50% of maximal inhibition was achieved, which was determined to be the p*K*_a_ of the catalytic cysteine residue. The p*K*_a_ of PsCHS and EaCHS are close to the 5.5 determined for other euphyllophyte CHSs, whereas the p*K*_a_ of PpCHS is over 1 pH unit higher, similar to that of SmCHS.

**Supplementary Figure 5.**
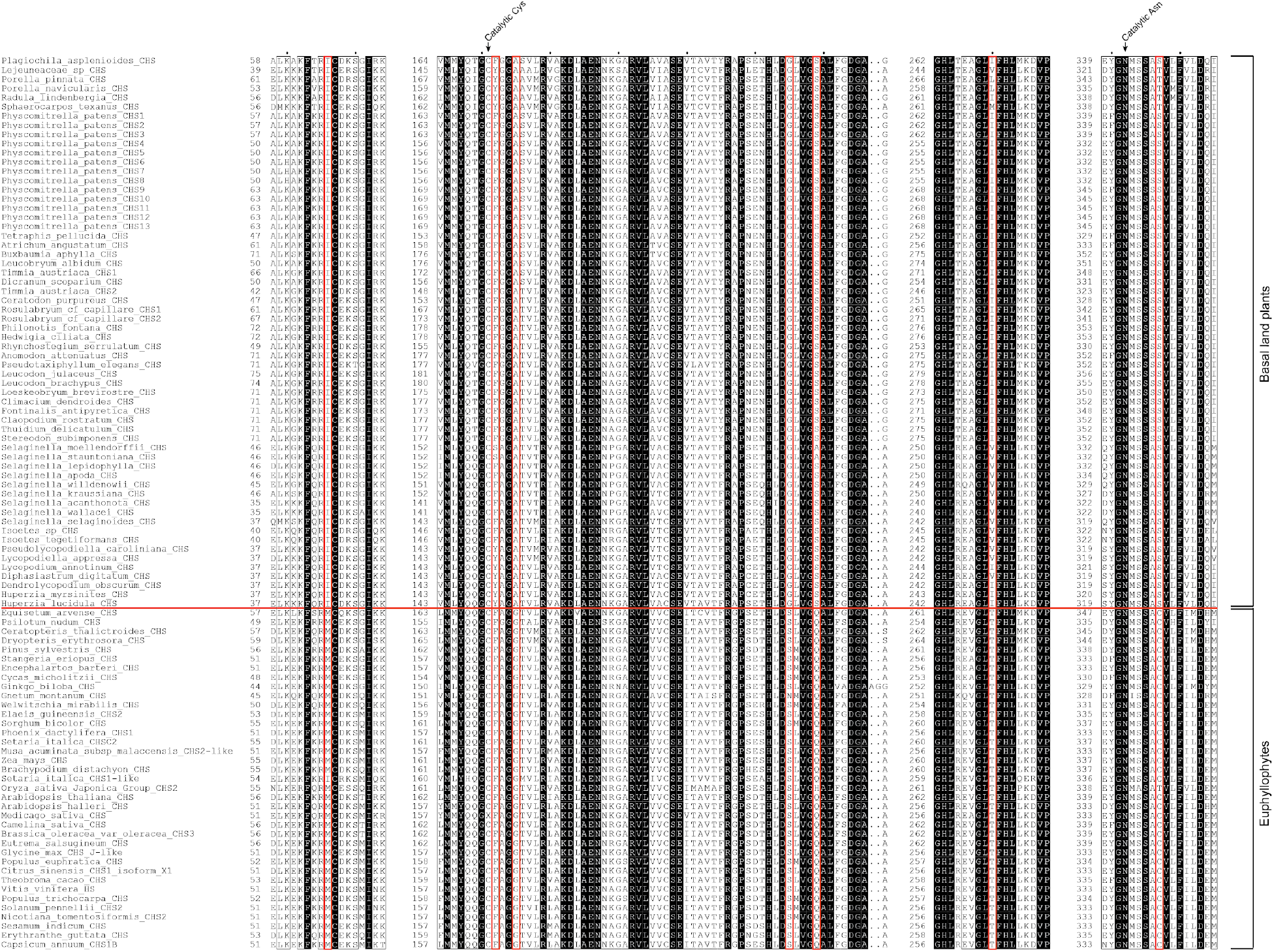
Multiple sequence alignment of CHSs. Sequence numbers of the beginning of each block for each CHS sequence are indicated. Residues outlined in thin black boxes are conserved with > 70% similarity across all sequences. Residues with 100% conservation are in white text with a black background. Red boxes indicate the seven positions mutated in the AtCHS M7 and SmCHS M7 constructs; these positions are differentially conserved between euphyllophyte and basal-plant CHSs, which are divided by the horizontal red line.

**Supplementary Figure 6.**
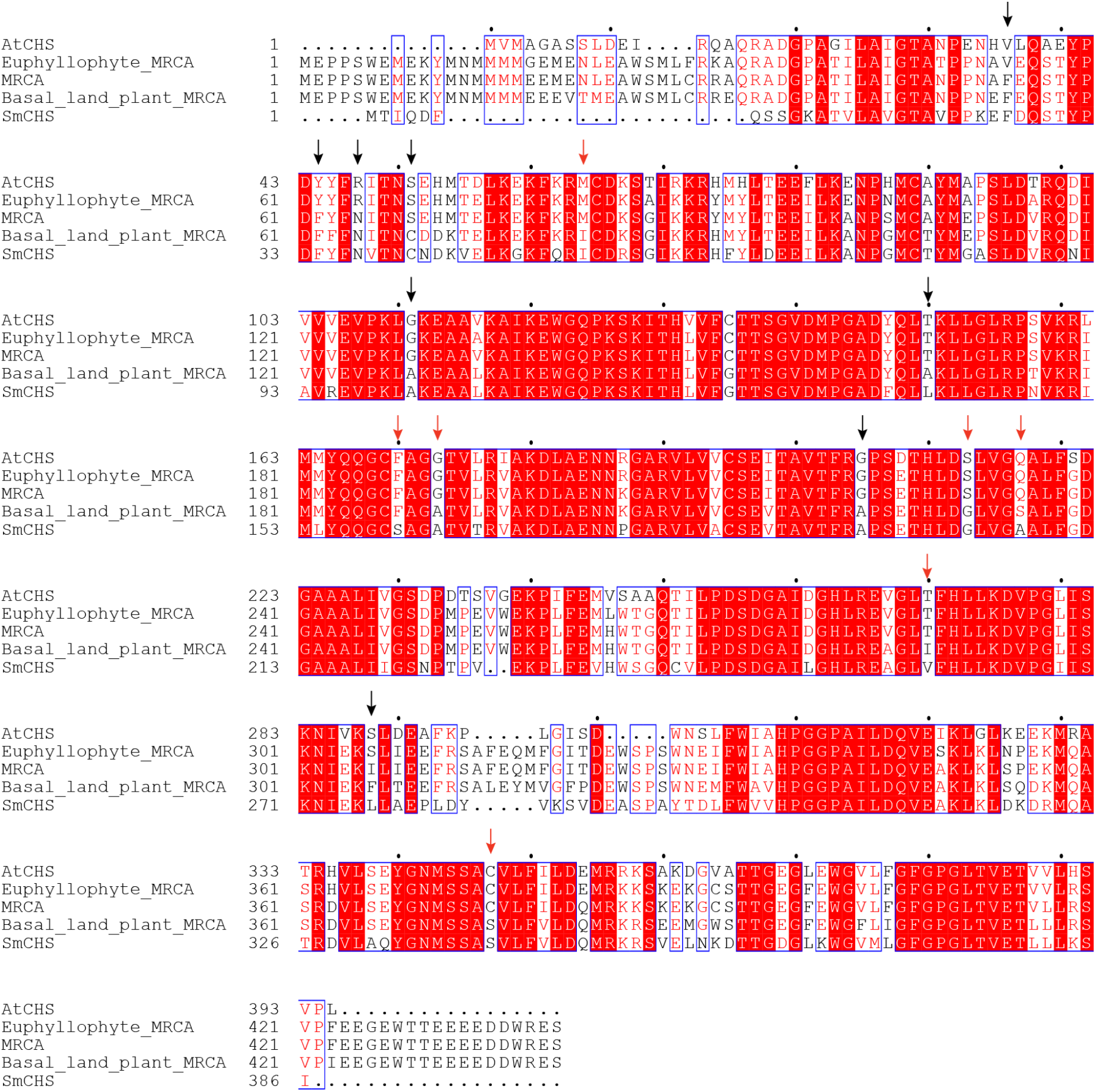
CHS ancestral sequence reconstruction. Sequences and phylogenetic tree of CHSs shown in Figure 1 were used to perform ancestral sequence construction with FastML. The most recent common ancestor (MRCA) sequences of all branches, euphyllophyte, and basal land plant clades are compared to AtCHS and SmCHS. Among the five sequences shown, absolutely conserved residues are shown in white text with red background. Residues with > 70% similarity are shown in red text and white background and blue outline. Other residues are shown in black text. Red arrows indicate the seven differentially conserved positions previously identified and mutated in the M7 CHS constructs. Black arrows indicate additional residue positions that are differentially conserved between euphyllophyte and basal-plant CHSs and determined to have possible functional impact based on their position in the CHS crystal structure.

**Supplementary Figure 7.**
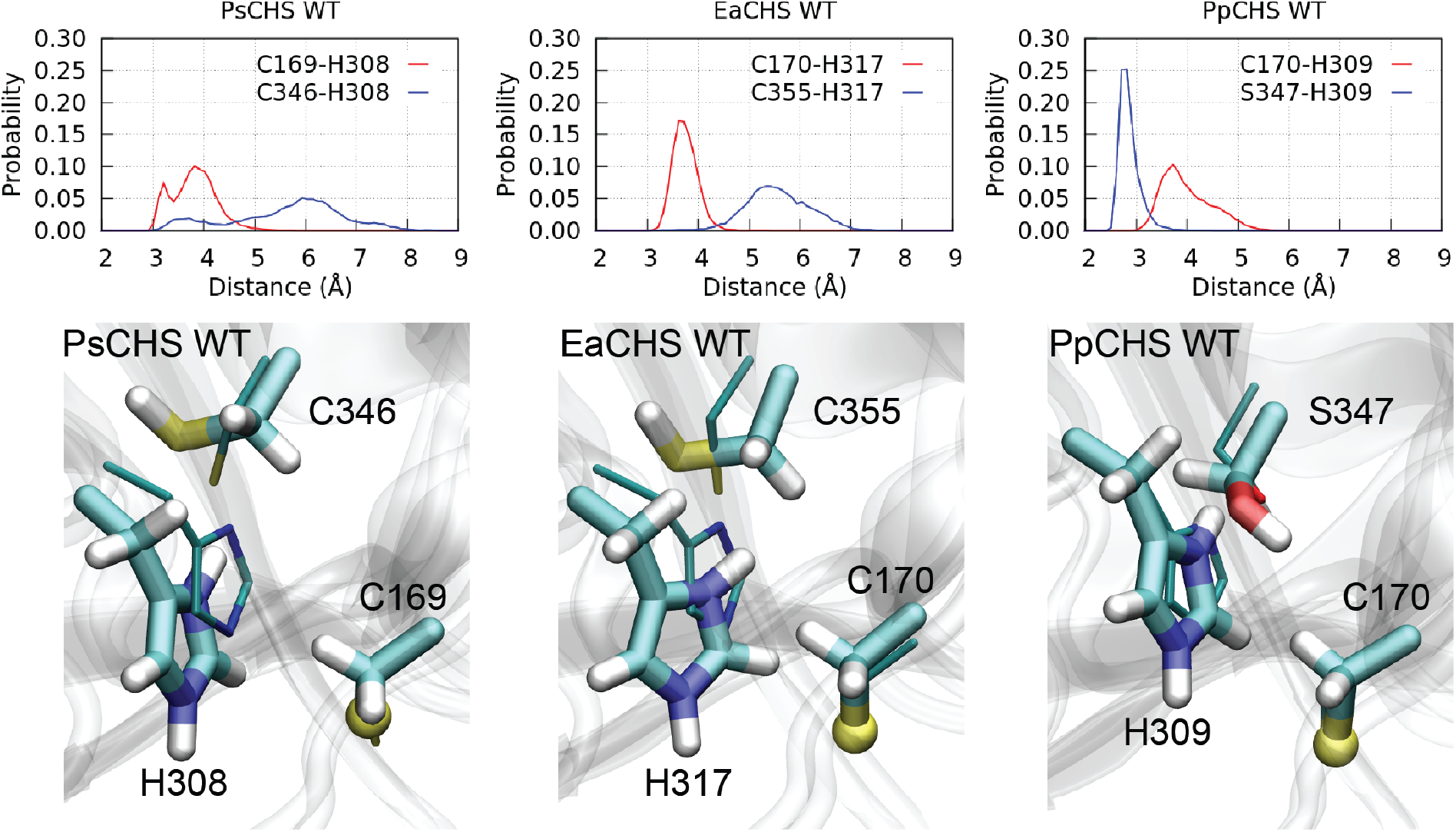
Distributions of inter-residue distances and the largest cluster conformations of EaCHS, PpCHS, PsCHS obtained from MD simulations. The observation of a serine forming a more stable hydrogen bond interaction than cysteine with the catalytic histidine is similar to the AtCHS and SmCHS wild-type and mutant simulations (Figure 5). Notably, with the rather weak interaction between the cysteine C346/C355 and the catalytic histidine, the latter moves more freely and often shows a much larger displacement from the corresponding position in the crystal structure (thin sticks).

**Supplementary Figure 8.**
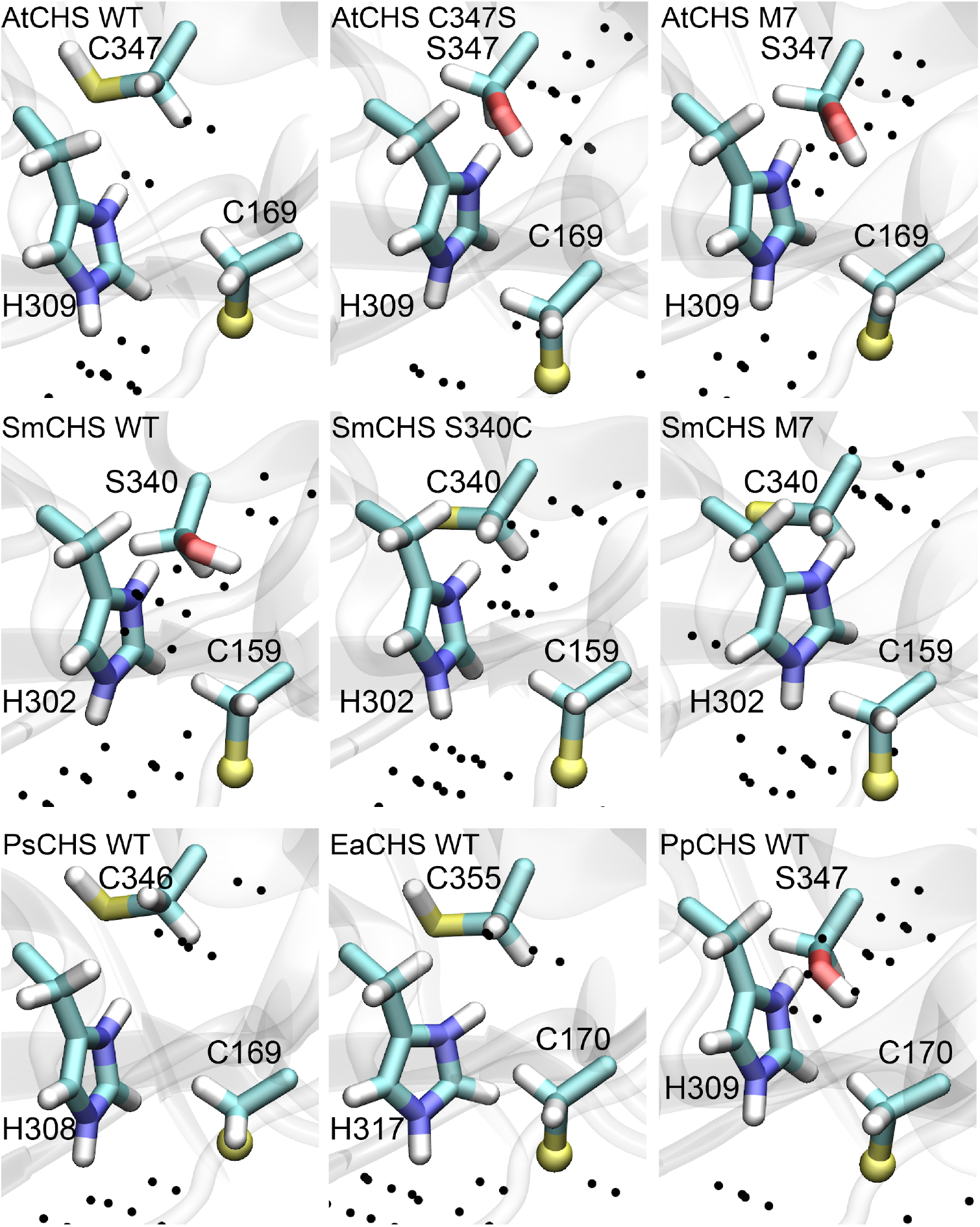
Average occupancy of water molecules obtained from MD simulations. Black dots represent grid points with an average water occupancy greater than 0.2. SmCHS in general has more water inside the active site, while the wild-type AtCHS has fewer water molecules. AtCHS mutants gradually attract more water around S347. This pattern is also observed in PpCHS, which also attracts more water around its serine than CHS where the serine is replaced by a cysteine (EaCHS, PsCHS).

**Supplementary Figure 9.**
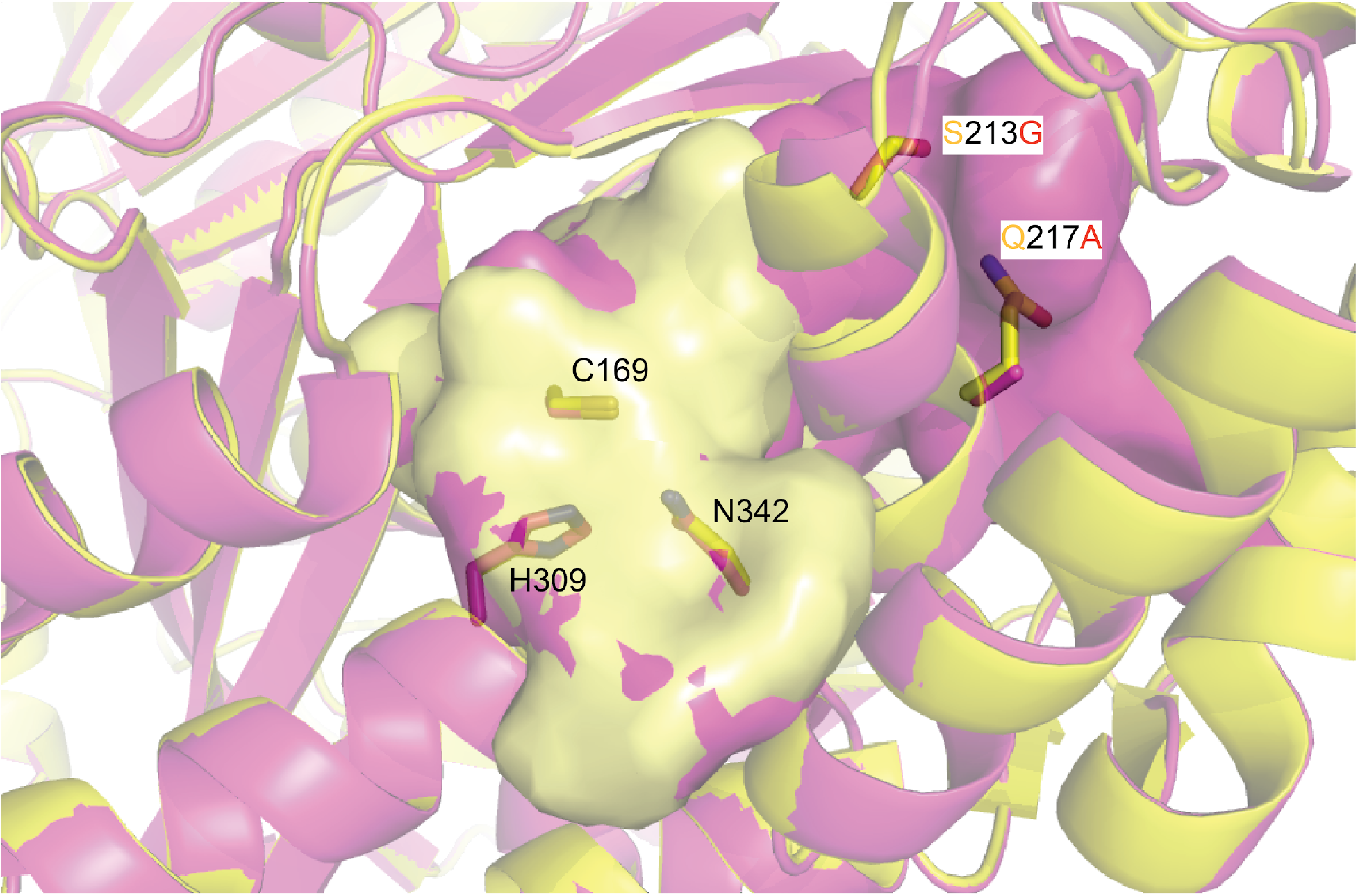
Comparison of wild-type AtCHS (yellow) and AtCHS M7 (yellow) crystal structures. The catalytic triad residues and two of the seven mutations from wild-type to M7 are modelled as sticks and labeled. The yellow and magenta surfaces represent the solvent-accessible cavities measured using the cavity-finding program KVFinder. The helix containing the two marked mutations is shifted in AtCHS M7 compared to wild type, leading to a larger active-site cavity.

